# Deficient ER Acetyl-CoA Import in Acinar Cells Leads to Chronic Pancreatitis

**DOI:** 10.1101/2020.03.25.008557

**Authors:** M. M. Cooley, D. D. H. Thomas, K. Deans, Y. Peng, A. Lugea, S. J. Pandol, L. Puglielli, G. E. Groblewski

**Affiliations:** Department of Nutritional Sciences, University of Wisconsin-Madison, Madison, WI, United States; Department of Medicine, University of Wisconsin-Madison, Madison, WI, United States; Pancreatic Research Group, Department of Medicine, Cedars-Sinai Medical Center, Los Angeles, CA, United States; Geriatric Research Education Clinical Center, Veterans Affairs Medical Center, Madison, WI, United States

## Abstract

Maintaining endoplasmic reticulum (ER) proteostasis is essential for pancreatic acinar cell function. Under conditions of severe ER stress, activation of pathogenic unfolded protein response pathways play a central role in the development and progression of pancreatitis. A key event in this pathogenic response is a loss of the transcription factor spliced XBP1 (XBP1s) and activation of the PERK pathway. Less is known of the consequence of perturbing ER-associated post-translational protein modification during pancreatitis. Here we show that expression of the ER acetyl-CoA transporter AT-1, necessary for ER protein acetylation, lies downstream of XBP1s and is significantly downregulated during the onset of pancreatitis. Genetic deletion of AT-1 in acinar cells of adult pancreas induces chronic ER stress marked by activation of both the XBP1s and PERK pathways, leading to mild/moderate chronic pancreatitis evidenced by accumulation of intracellular trypsin, immune cell infiltration, and fibrosis, but little pancreatic degeneration. Two-day induction of acute on chronic pancreatitis in AT-1 acinar specific knockout mice results in a severe CP phenotype with pronounced pancreatic atrophy. These findings uncover a new layer of complexity of the pathological ER stress response and its impact on pancreatic disease.

## Introduction

Chronic pancreatitis (CP) is a persistent fibroinflammatory condition that to date lacks specific clinical therapies (1). The mechanisms of CP pathogenesis and progression remain incompletely understood as they involve a complex interplay of genetic and environmental factors. One cellular event consistent across nearly all experimental models of both acute (AP) and CP is the pathological activation of the unfolded protein response (UPR), and thus continues to be a pathway of interest in the study of pancreatic diseases (2–4).

UPR activity is critical and necessary to maintain pancreatic acinar cell secretory function. Importantly, XBP1s acts as a terminal differentiation transcription factor in concert with Mist1 in exocrine cells (5, 6). Previous studies have demonstrated that IRE1/XBP1s activates an adaptive response to endoplasmic reticulum (ER) stress and protects acinar cells from damage (7–9). On the other hand, PERK signaling in the face of unresolved ER stress leads to upregulation of cell death regulator CHOP via ATF4 (10, 11). The resulting acinar cell injury and death induces pancreatitis responses including initiation of an inflammatory cascade and, in CP pathology, stellate cell activation and concomitant collagen deposition and fibrosis. Consistent with these data, it was recently shown that pancreatitis stimuli in human acinar cells reduces protective XBP1s signaling and in turn upregulates CHOP, cell damage, and activation of inflammatory signaling (12). In the past several years, mutations associated with digestive enzyme misfolding have been identified that, when introduced in mouse models, induce ER stress responses and exhibit spontaneous CP (13–17).

ER proteostasis is influenced in part by Nε-lysine acetylation of nascent proteins, a post-translational modification that assists in protein stability and trafficking through the secretory pathway (18). This process is regulated by the ER acetyl-CoA transporter AT-1, which moves cytosolic acetyl-CoA into the ER lumen to be used as a substrate for resident acetyltransferases (19, 20). Disruption of AT-1 activity in both humans and mice results in various neurological defects involving chronic ER stress responses, secretory efficiency, and autophagy regulation. In the human population, a S113R mutation in AT-1 manifests as hereditary spastic paraplegias, and AT-1^S113R/+^ hypomorphic mice exhibit both neurodegeneration and impaired immune surveillance (21). Chromosomal duplications that include the AT-1 gene are associated with cognitive dysfunction and dysmorphism in humans, and overexpression of AT-1 in mouse models yields a progeria-like phenotype (22, 23). These studies underscore the importance of AT-1 activity on protein acetylation and maintenance of ER proteostasis.

Given the robust systemic phenotypes seen with altered AT-1 expression, we theorized that the same chronic UPR activation would likely translate into pancreatic pathophysiology. Here, we investigated the effects of chronic ER stress in AT-1^S113R/+^ hypomorphic mice and inducible acinar specific AT-1 knockout mice on pancreatic function. Our findings support that disruption of AT-1 activity in pancreas leads to persistent UPR activity, inflammation, and fibrosis consistent with mild/moderate CP that progresses to severe disease with limited recovery when challenged with AP.

## Results

### Relationship of AT-1 expression, ER stress, and pancreatitis

It was previously reported that AT-1 expression is regulated downstream of the IRE1/XBP1s pathway (24). Consistent with this, AT-1 mRNA is significantly downregulated in acinar-specific Ela-Cre XBP1^−/−^ pancreas (Figure 1A), which display spontaneous pancreatic disease (unpublished results). XBP1s is upregulated in acinar cells during AP as a protective mechanism against ER stress-induced damage, then significantly decreases as the disease progresses (12). Accordingly, AT-1 mRNA is reduced by greater than 50% following cerulein induced AP (CER-AP) in WT mice (Figure 1B, left), while AT-1 mRNA was decreased in 75% of WT mice subject to CER-CP (Figure 1B, right). These results suggest that AT-1 loss during pancreatitis is secondary to the reduction in XBP1s activity and, based on the current results, likely exacerbates disease progression.

**Figure 1.**
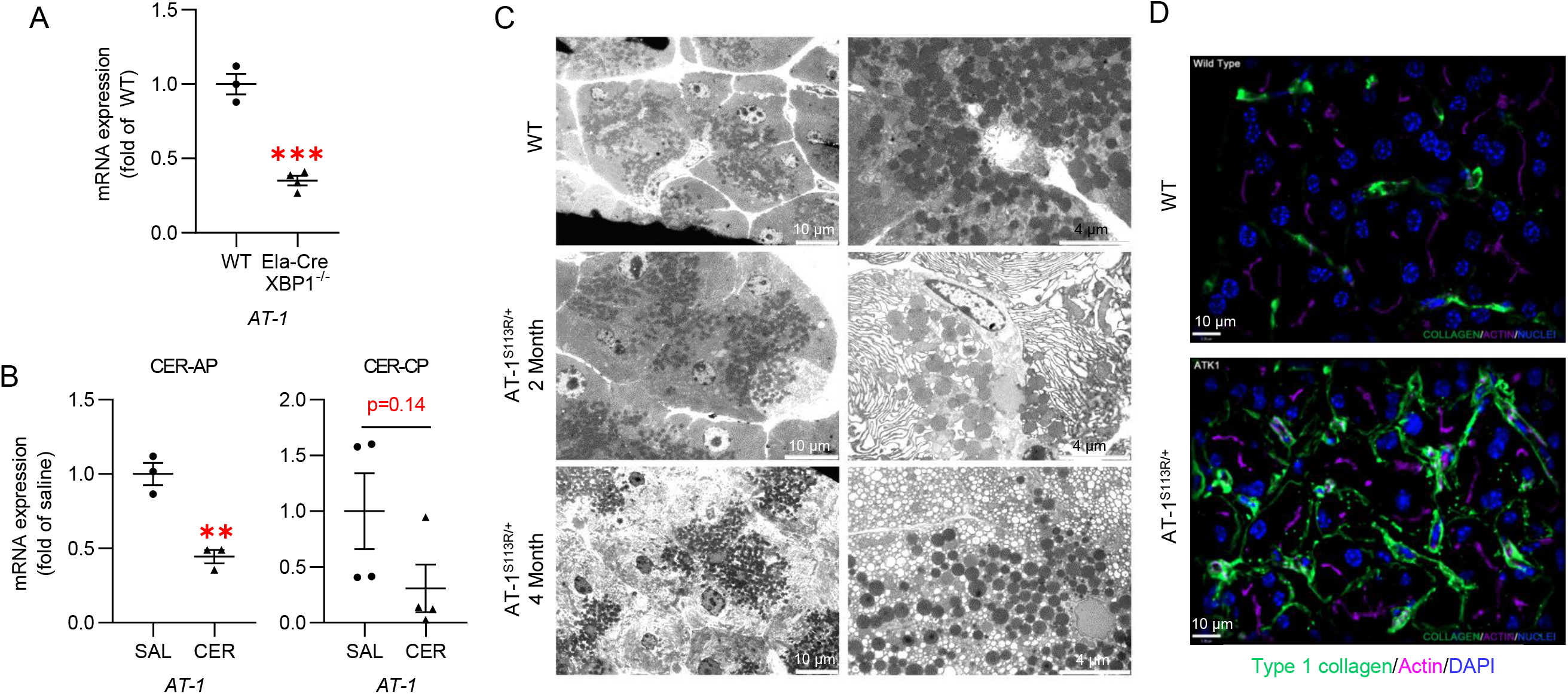
AT-1 reduction is associated with pancreatitis. **A.** AT-1 RNA expression in pancreas from WT or Ela-Cre XBP1^−/−^ mice. **B.** AT-1 mRNA expression in pancreas from WT mice treated with saline (SAL) or cerulein (CER) to induce acute pancreatitis (CER-AP, left panel) or chronic pancreatitis (CER-CP, right panel). **C.** Electron microscopy of pancreatic tissue from WT and hypomorphic AT-1^S113R/+^ pancreas at 2 and 4 months of age. Scale bars: 10μm (left panels) and 4μm (right panels) **D.** Immunofluorescence of type 1 collagen (green) and actin (purple) in WT and AT-1^S113R/+^ pancreatic sections. Scale bars: 10 μm. Data are mean +/− SE, n=3-5, ***p<0.001, ****p<0.0001. Statistical comparison of means was done by unpaired two-tailed Student’s *t*-test.

Given the critical role of AT-1 in maintaining ER proteostasis and the well-defined relationship of elevated ER stress and pancreatitis, we examined whether dysregulation of AT-1 function itself would be sufficient to induce pancreatitis. AT-1^S113R/+^ mice, which express a hypomorphic mutation that prevents AT-1 dimerization, have an approximate 50% reduction in ER acetyl-CoA transport activity (S113R homozygosity is embryonic lethal) (21). Notwithstanding the immune dysfunction reported in these mice, AT-1^S113R/+^ acinar cells display widespread ER dilation and vacuolization, indicative of ER stress and cellular damage, which progresses over time (Figure 1C). Likewise, AT-1^S113R/+^ pancreas have increased collagen deposition (Figure 1D), a marker of fibrosis.

### Generation of acinar-specific AT-1 KO mice

To circumvent immune system interference, we crossed AT-1 floxed (Slc33a1^tm1a(KOMP)Wtsi^) mice with mice expressing tamoxifen-inducible Cre driven by the acinar-specific elastase promoter (Tg(Cela1-cre/ERT)1Lgdn/Jd (25)) to produce acinar-specific heterozygous and homozygous AT-1 KO’s (Ela-Cre AT-1^+/−^ and Ela-Cre AT-1^−/−^, respectively). To allow for normal pancreas development, tamoxifen was administered by oral gavage at 2 months of age and tissues were analyzed 2-3 months post tamoxifen (Figure 2A). Pancreatic AT-1 mRNA was reduced by half in Ela-Cre AT-1^+/−^ and approximately 90% loss in Ela-Cre AT-1^−/−^ (Figure 2B), with no changes observed in liver (Figure 2C). As AT-1 mRNA was measured from whole pancreatic tissue, residual AT-1 transcript expression in Ela-Cre AT-1^−/−^ is from cells other than acini present in the mixed gland. No significant changes in body weight or pancreatic weight relative to body weight were observed at 4-5 months of age (Figure 2D) out to 1.5 years (data not shown). No overt differences in histology or biochemical parameters were seen in Ela-Cre mice given either tamoxifen or corn oil alone (Supplementary Figure 1A-C).

**Figure 2.**
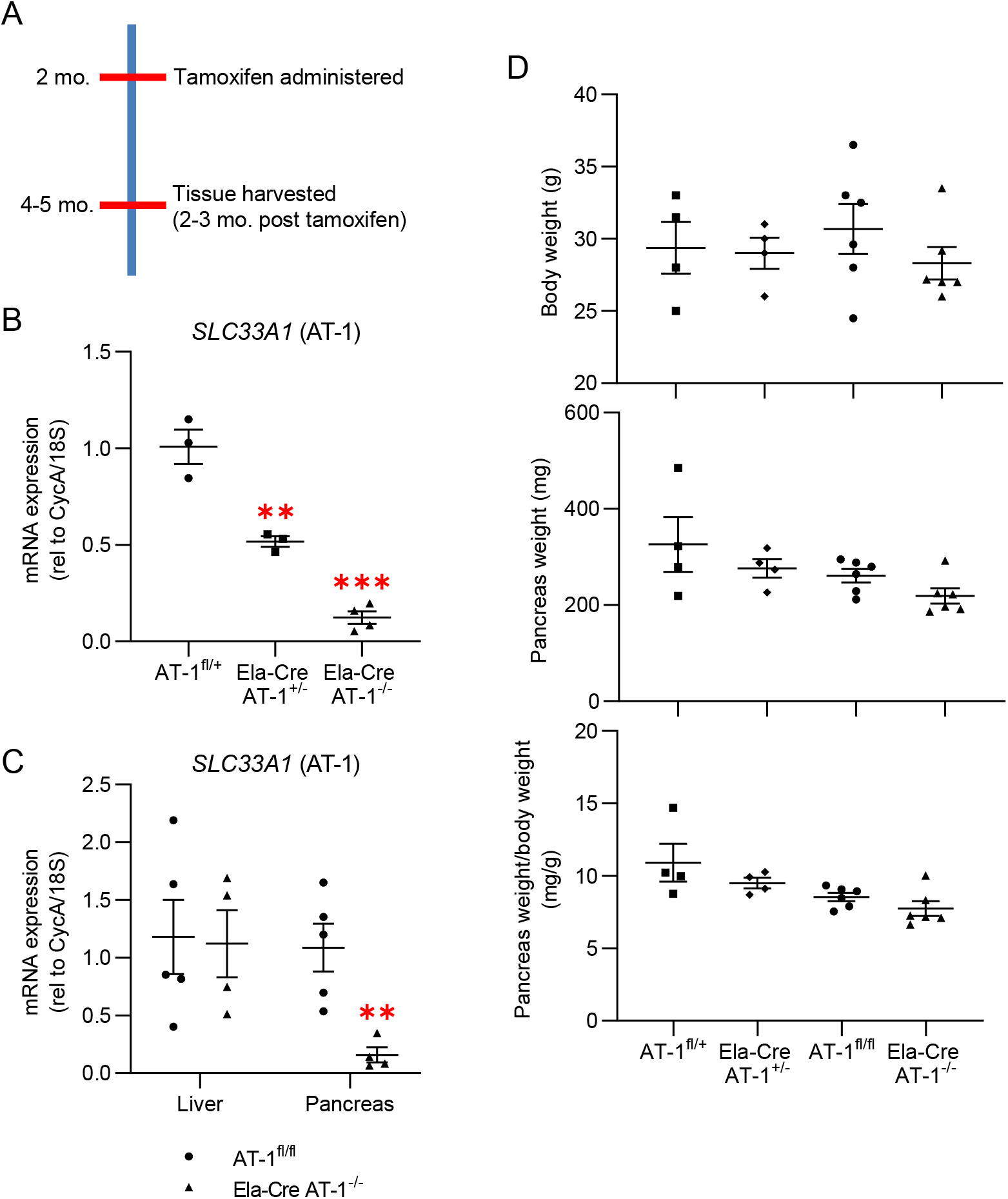
Characterization of tamoxifen inducible acinar-specific AT-1 mice. **A.** Schematic of Ela-Cre AT-1 mouse production. KO was induced by tamoxifen at 2 months of age to allow for normal pancreatic development; tissues were analyzed 2-3 months post. **B-C** *SLC33A1* (AT-1) levels in whole pancreas and/or liver from AT-1^fl/+^, Ela-Cre AT-1^+/−^, and Ela-Cre AT-1^−/−^ mice, as indicated. **D.** Measurement of body weight, pancreas weight, and pancreas weight normalized to body weight between AT-1^fl/+^, Ela-Cre AT-1^+/−^, AT-1^fl/fl^, and Ela-Cre AT-1^−/−^ mice. Data are mean +/− SE, n=3-5, **p<0.01, ***p<0.001. Statistical comparison of means was done by unpaired two-tailed Student’s *t*-test (C) or one-way ANOVA (B, D).

### AT-1 deletion induces chronic ER stress

Electron microscopy of Ela-Cre AT-1^−/−^ acini revealed ER dilation and accumulation of vacuoles, recapitulating the phenotype observed in AT-1^S113R/+^ in Figure 1 (Figure 3A). Extensive UPR activation occurred marked by an 8-fold increase in XBP1s protein (Figure 3B, C) and a reduction in total XBP1 mRNA (Figure 3D). Furthermore, JNK activation was significantly increased in Ela-Cre AT-1^−/−^, likely due to IRE1 hyperstimulation (Figure 3E, F) (26–28). ATF6 mRNA was also elevated in Ela-Cre AT-1^−/−^ compared to controls (Figure 3G). Finally, PERK and its downstream targets peIF2α, ATF4, GADD34, and cell death mediators CHOP and cleaved caspase-3 were all significantly increased (Figure 3H, I, J); notably, CHOP protein expression was increased 4-fold and mRNA expression increased approximately 10-fold (Figure 3I, J). Collectively, these results indicate that loss of ER acetyl-CoA availability via AT-1 deletion activates a pronounced response involving all known branches of the ER stress pathway.

**Figure 3.**
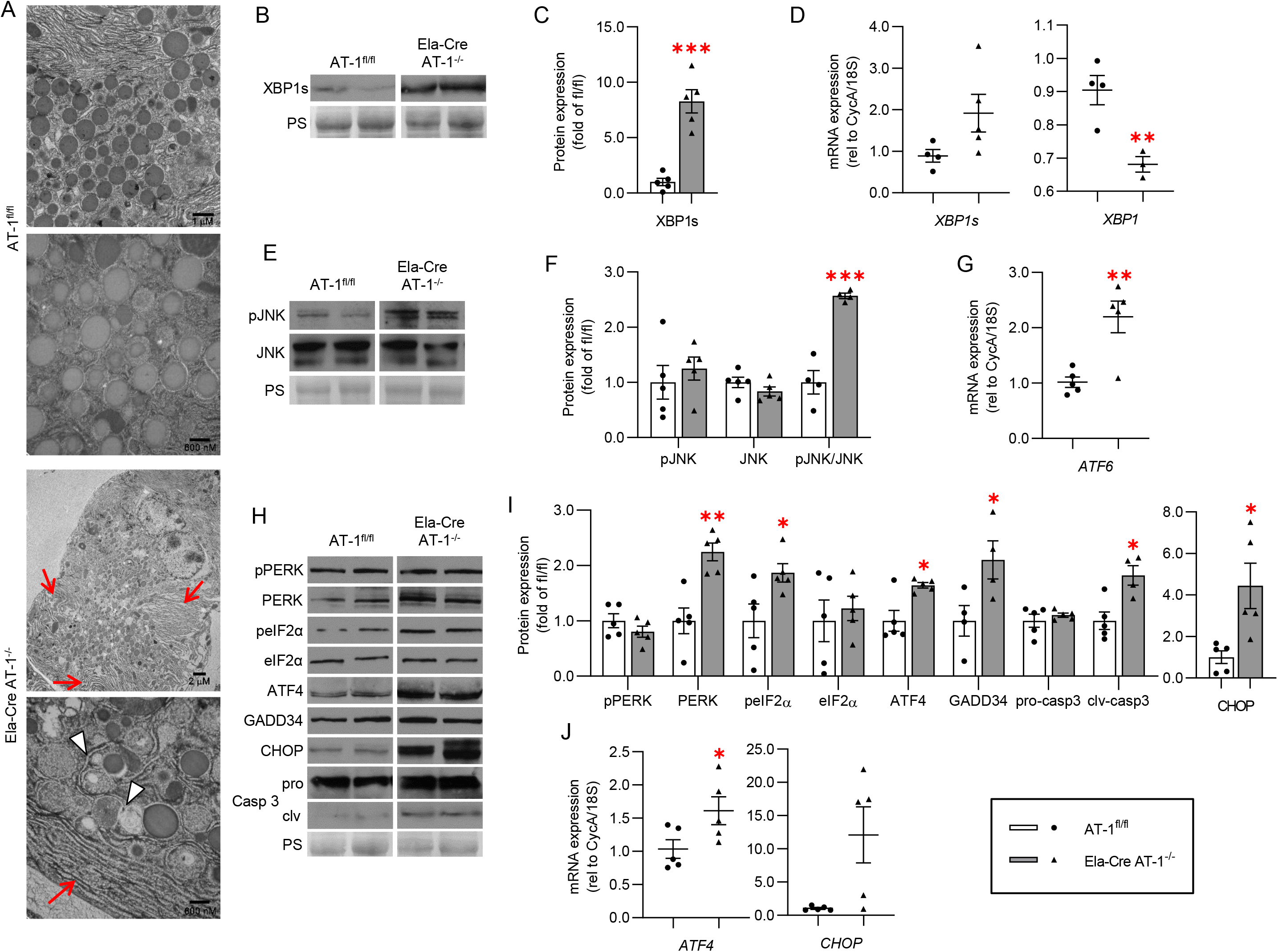
Loss of AT-1 induces ER stress in pancreas. **A.** Representative EM images of pancreatic sections from AT-1^fl/fl^ and Ela-Cre AT-1^−/−^. Red arrows indicate dilated ER; white arrowheads indicate vacuoles. Scale bars: 2μm (top panel) and 800nm (bottom) in AT-1^fl/fl^; 1μm (top) and 600nm (bottom) in Ela-Cre AT-1^−/−^. **B.** Immunoblot of sXBP1, quantified in **C**. **D.** qPCR assessment of *XBP1s*, and *XBP1 (tot)* mRNA. **E.** Immunoblot of phosphorylated and total JNK, quantified in **F**. **G.** qPCR of *ATF6*. **H.** Immunoblot of pPERK, PERK, peIF2α, eIF2α, ATF4, GADD34, caspase 3 (pro- and cleaved forms), and CHOP, quantified in **I**. **J.** qPCR analysis of *ATF4* and *CHOP* mRNA. Immunoblot and qPCR were done in whole pancreas. Ponceau S (PS) is loading control. All data are mean +/− SE, n=4-5, *p<0.05, **p<0.01, ***p<0.001, ****p<0.0001. Statistical comparison of means was done by unpaired two-tailed Student’s *t*-test.

### Ela-Cre AT-1^−/−^ pancreas exhibits inflammation and fibrosis consistent with CP

Robust immune cell infiltration was evident by H&E staining in Ela-Cre AT-1^−/−^ pancreas, a key pathological event in pancreatitis (Figure 4A). Congruent with this observation, we found significant NF-κB activation (Figure 4B, C) and a 5-fold increase in expression of the potent chemokine Ccl5 (Figure 4D), a target of NF-κB. Immune cell species present in Ela-Cre AT-1^−/−^ pancreas include predominantly F4/80^+^ macrophages and, to a lesser extent, CD45R^+^ B cells and CD3^+^ T cells (Figure 4E). Furthermore, Ela-Cre AT-1^−/−^ pancreas exhibited significant α-smooth muscle actin upregulation and collagen deposition by Sirius Red staining, indicative of fibrosis (Figure 5A, B, C). Taken together, the presence of persistent UPR activation, autophagic defects, chronic inflammation, and fibrosis support a CP phenotype with AT-1 deletion.

**Figure 4.**
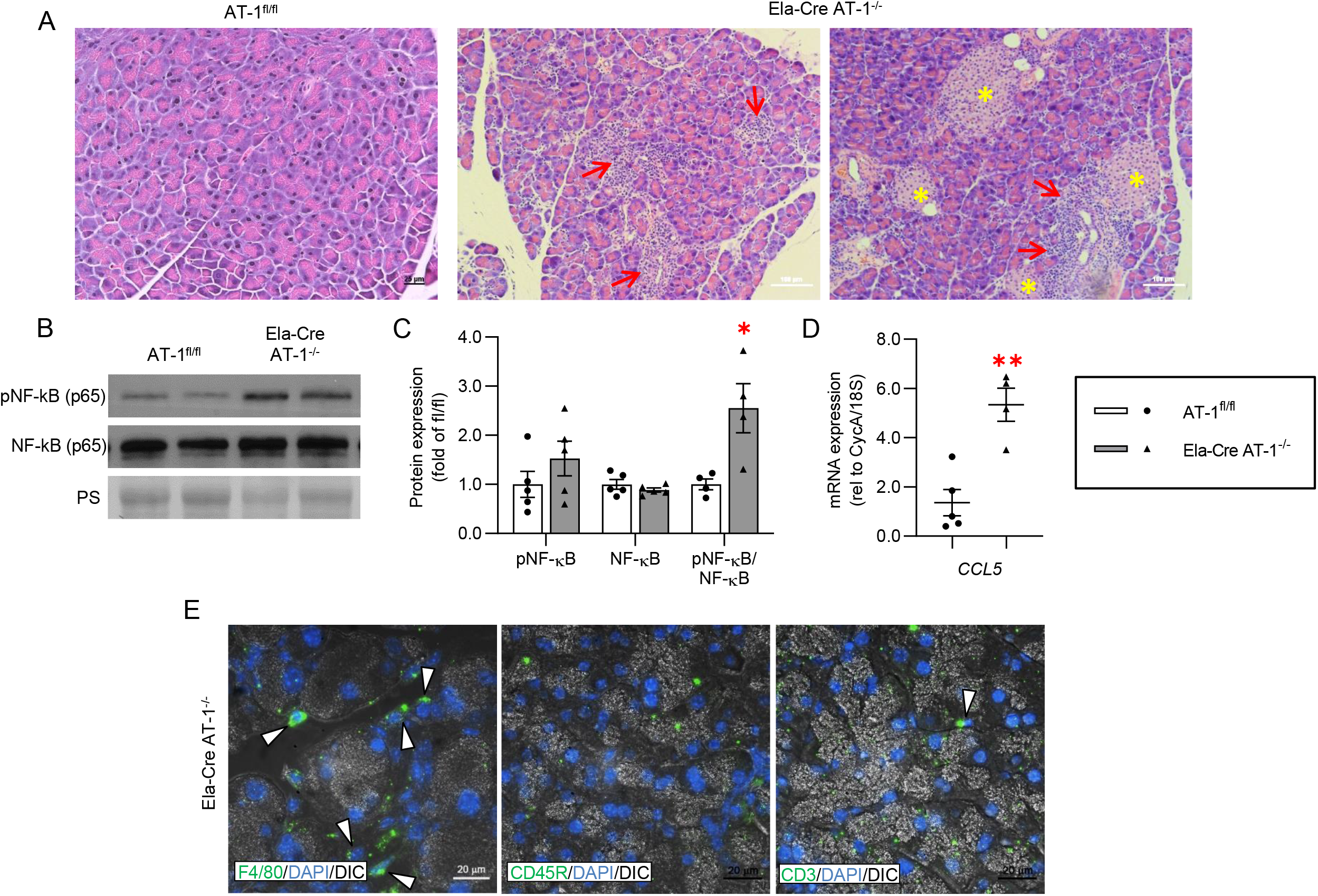
Loss of AT-1 results in inflammation. **A.** Representative H&E stainings of AT-1^fl/+^ and Ela-Cre AT-1^−/−^ pancreatic sections. Note significant infiltration of immune cells in Ela-Cre AT-1^−/−^ (red arrows). Islets are marked with yellow asterisks. Scale bars: 25 (left panel) and 100μm (middle and right panels). **B.** Immunoblot of phosphorylated and total NF-κB (p65), quantified in **C**. Ponceau S (PS) is loading control. **D.** qPCR analysis of *CCL5*. **E.** Representative IF images of immune cell markers F4/80, CD45R, and CD3 (white arrowheads). Scale bars: 20μm. Immunoblot and qPCR was done in whole pancreas. All data are mean +/− SE, n4-5, *p<0.05, **p<0.01. Statistical comparison of means was done by unpaired two-tailed Student’s *t*-test.

**Figure 5.**
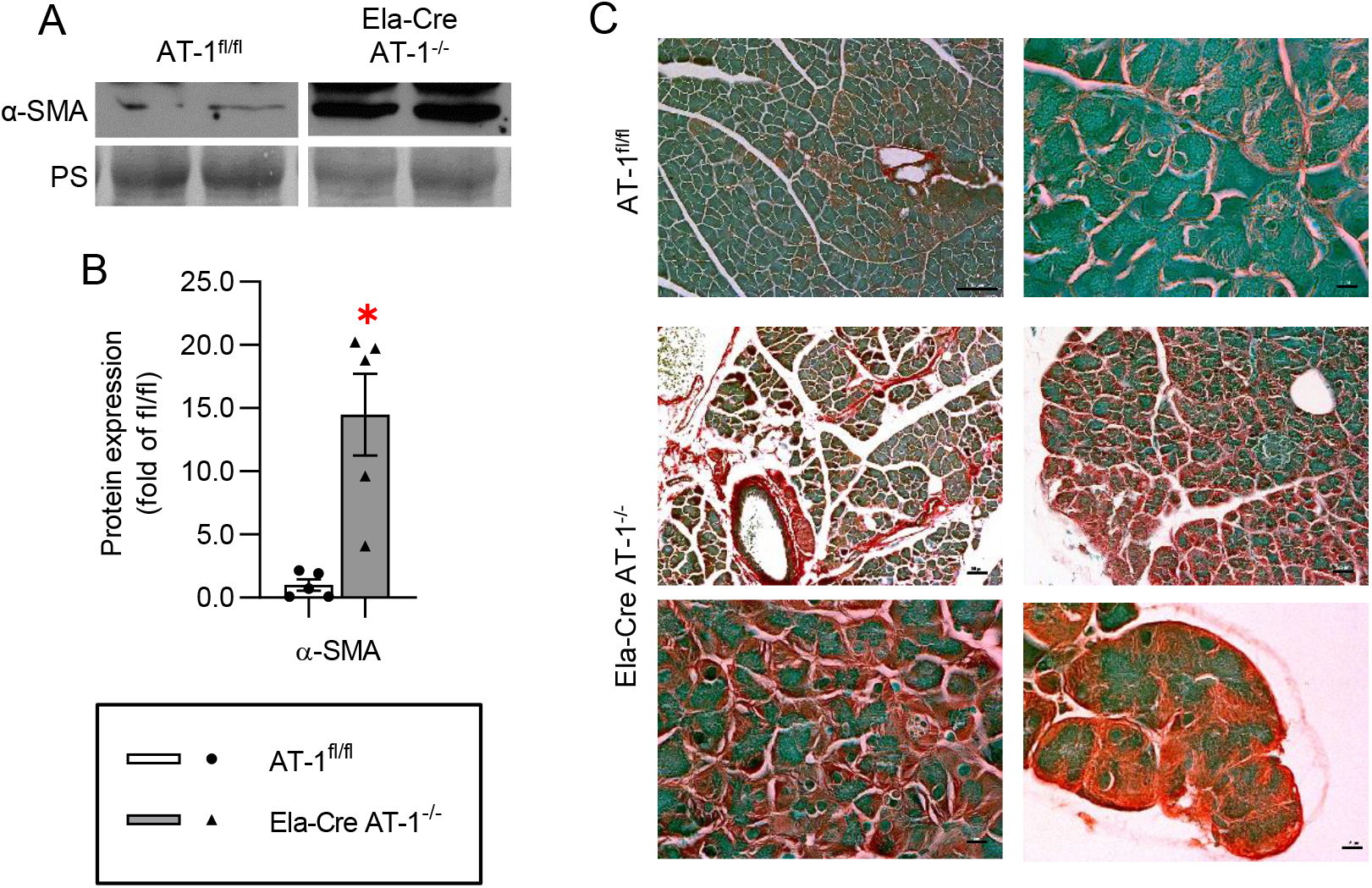
AT-1 deletion induces fibrosis. **A.** Immunoblot of alpha smooth muscle actin (α-SMA) in AT-1^fl/fl^ and Ela-Cre AT-1^−/−^ whole pancreas, quantified in **B**. Ponceau S (PS) is loading control. **C.** Representative Sirius red staining of AT-1^fl/fl^ and Ela-Cre AT-1^−/−^ pancreatic sections. Scale bars: (from top left to bottom right) 100, 10, 50, 25, 10, and 10μm. Data are mean +/− SE, n=4-5, *p<0.05. Statistical comparison of means was done by unpaired two-tailed Student’s *t*-test.

### Anomalies in digestive enzyme expression, amylase secretion, and trypsin activation

Examination of digestive enzyme content revealed a more than two-fold increase in amylase yet significant decreases in lipase and elastase expression in Ela-Cre AT-1^−/−^ (Figure 6A, B, C). Despite this ostensible intracellular accumulation of amylase, amylase secretion from isolated acini was significantly elevated between 3-30pM CCK-8 in Ela-Cre AT-1^−/−^ compared to AT-1^fl/fl^ controls when either normalized as a percent of total amylase (data not shown) or to cellular DNA (Figure 6D). Unexpectedly, enhanced trypsin activation under basal conditions was detected in Ela-Cre AT-1^−/−^ pancreas, a less common phenomenon seen in CP vs AP (Figure 6E). No differences in plasma amylase were observed (Figure 6F).

**Figure 6.**
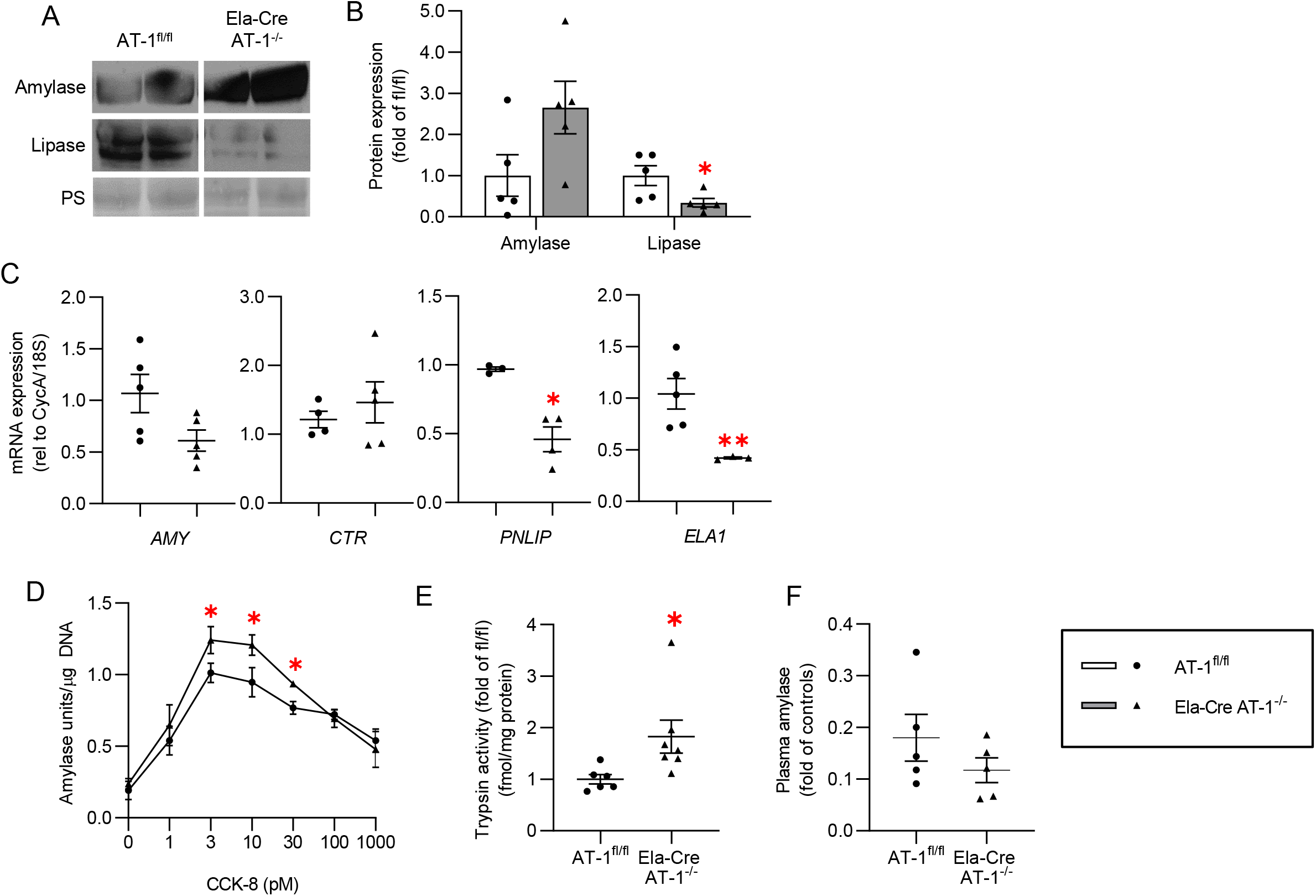
Digestive enzyme content and release is altered with AT-1 deletion. **A.** Immunoblot of amylase and lipase, quantified in **B.** Ponceau S (PS) is loading control. **C.** qPCR assessment of digestive enzyme mRNA expression in whole pancreas of AT-1^fl/fl^ and Ela-Cre AT-1^−/−^ mice. **D.** Amylase release normalized to cellular DNA from isolated acinar cells stimulated with increasing amounts of CCK-8. **E.** Plasma amylase activity in AT-1^fl/+^, Ela-Cre AT-1^+/−^, AT-1^fl/fl^, and Ela-Cre AT-1^−/−^. **F.** Measurement of trypsin activation. Data are mean +/− SE, n≥3, *p<0.05, **p<0.01. Statistical comparison of means was done by unpaired two-tailed Student’s *t*-test.

### Acute on chronic pancreatitis increases disease severity in Ela-Cre AT-1^−/−^

Despite the chronic ER stress, inflammation, and fibrosis, no pancreatic degeneration as noted by pancreatic weight was found with AT-1 deletion (see Figure 2D), suggesting a mild/moderate CP phenotype. To examine the response to AP challenge, AT-1^fl/fl^ and Ela-Cre AT-1^−/−^ mice were subject to *in vivo* CER-AP for 2 consecutive days then analyzed following a 1-week recovery period (Figure 7A). While AT-1^fl/fl^ control mice showed pancreatic recovery, Ela-Cre AT-1^−/−^ pancreas underwent significant atrophy with a 40% reduction in gland weight compared to controls, but no change in body weight (Figure 7B, C). This was confirmed by H&E and Sirius Red staining showing extensive acinar cell loss, inflammatory cell infiltration, fat replacement, and fibrosis (Figure 7D, E). Together, these results support that inducing acute on CP in Ela-Cre AT-1^−/−^ results in a severe CP phenotype.

**Figure 7.**
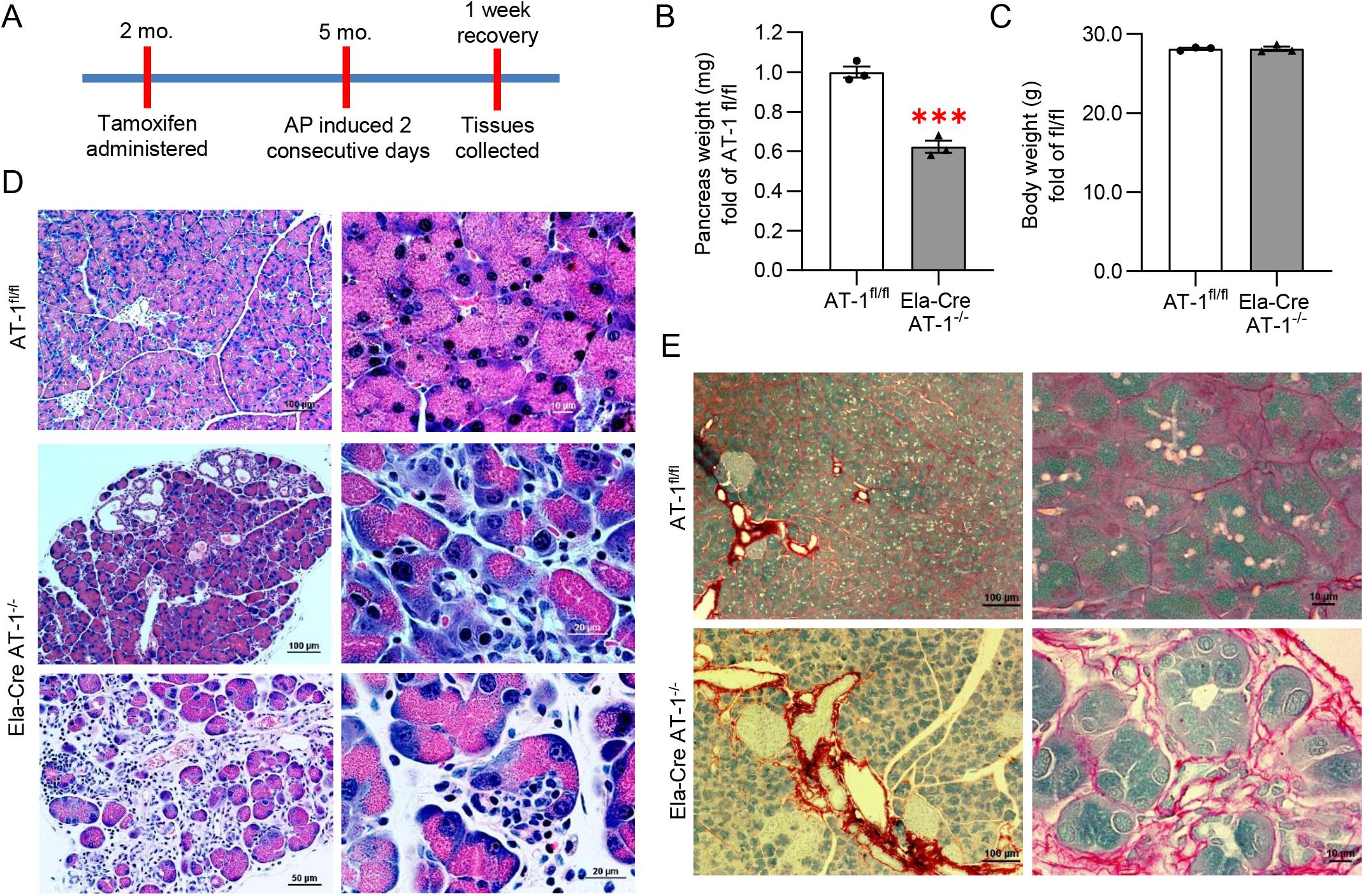
Ela-Cre AT-1^−/−^ sustain significant damage during acute pancreatitis. **A.** Experimental schematic: AT-1^fl/fl^ and Ela-Cre AT-1^−/−^ mice given CER-AP and allowed to recover 1 week. **B-C.** Pancreatic weight (B) and body weight (C) post-experiment. **D.** Representative H&E staining of AT-1^fl/fl^ and Ela-Cre AT-1^−/−^ pancreatic sections. Scale bars: 100μm (left) and 10μm (bottom) for AT-1^fl/fl^; 100μm (top left), 50μm (bottom left), and 20μm (right panels) for Ela-Cre AT-1^−/−^. **E.** Representative images of collagen Sirius red staining. Scale bars: 100μm (left panels) and 10μm (right panels). Data are mean +/− SE, n=3, ***p<0.001. Statistical comparison of means was done by unpaired two-tailed Student’s *t*-test.

## Discussion

Pathological UPR activation is a key cellular event in AP development, and chronic UPR activity occurs in models of CP (3). Here, we demonstrate that chronic ER stress induced by the loss of the ER acetyl-CoA transporter AT-1 in pancreatic acinar cells is sufficient to cause spontaneous mild/moderate CP. Deletion of AT-1 leads to broad and pronounced UPR upregulation, inflammation, fibrosis, and, in response to CER-AP, significant gland atrophy and acinar cell loss.

Previous studies targeting ER stress regulatory molecules have shed light on their effects on pancreatic function and disease pathology. Pancreas from PERK^−/−^mice, in spite of acinar ER morphological abnormalities, develop normally and are protected from further damage by compensation from IRE1/XBP1s (29). In contrast, acini lacking XBP1 undergo apoptosis during embryogenesis, revealing the critical role XBP1 plays in acinar cell development (5). Interestingly, XBP1^+/−^ mice display normal pancreatic function until challenged with alcohol, supporting that enhanced XBP1 expression is protective against pancreatitis onset (9). The present study demonstrates a marked upregulation and/or dysregulation of pathological cellular pathways including PERK, CHOP, and JNK with loss of AT-1; however, the CP observed in Ela-Cre AT-1^−/−^ appeared to be relatively mild to moderate, suggesting a capability of acinar cells to adapt to chronic ER stress when the UPR is intact. Additionally, a recent study suggests that chronic ER stress induces a switch to eIF3-dependent translation, circumventing eIF2α inhibition and partially restoring protein translation including that of ER functional proteins (30). These compensatory mechanisms allow acini to continue functioning under high stress conditions and may explain, in part, how some cases of CP in humans are asymptomatic (31); indeed, CP manifestation in patients with high alcohol intake varies significantly and, in some cases, is only detected postmortem (32, 33). Furthermore, severe CP due to alcohol intake is thought to occur following recurrent bouts of clinical or subclinical AP (34). Similarly, AT-1 knockout mice progressed to severe CP with loss of pancreatic mass only after repeated CER-AP. These results support the concept that the combination of environmental stressors and subclinical pathophysiological perturbations supersede the adaptive capacity of the UPR in acinar cells and promote severe disease.

The effect of AT-1 deletion on pancreatic function contributes to a growing understanding of the significance of Nε-lysine acetylation in the ER on protein folding and ER stress responses (18, 35, 36). Investigation of the ER acetylome in cultured cell models identified proteins such as BiP, LAMP2, and cathepsin D as acetylation targets (37), all of which are critical for pancreatic acinar cell function (38–41). Further, studies have shown increased UPR activity and CP-like outcomes in response to variants of the digestive enzymes *PRSS1* and *CPA1* among others that lead to their misfolding (15). The unexpected finding of increased trypsin activation as well as variations in the expression of other digestive enzymes with AT-1 deletion calls into question the acetylation state of these proteins and what the functional significance of these posttranslational modifications might be on pancreatitis outcomes. Further investigation into the pancreatic acinar cell ER acetylome will be necessary to reveal additional insights.

## Experimental Methods

### Reagents

#### Antibodies

Antibodies utilized for immunoblotting and/or immunofluorescence and their source are indicated in Table 1 below. Peroxidase-conjugated secondary antibodies were from GE Healthcare. Alexa-conjugated secondary antibodies were from Life Technologies.

**Table 1.** Primary antibodies.

| **Target** | **Source** | **Cat #** |
| --- | --- | --- |
| Actin | AbCam | Ab6276 |
| Amylase | Sigma-Aldrich | A827 |
| ATF4 | Santa Cruz | sc-200 |
| Caspase 3 | Thermo Fisher | PA5-16385 |
| CD3 | AbCam | Ab16669 |
| CD45R | AbCam | Ab64100 |
| CHOP | Cell Signaling | cs-2895 |
| eIF2α | Cell Signaling | cs-9722 |
| p-eIF2α | Cell Signaling | cs-9721 |
| F4/80 | AbCam | Ab6640 |
| GADD34 | Santa Cruz | sc-8327 |
| JNK | Cell Signaling | cs-9252 |
| p-JNK | Cell Signaling | cs-4668 |
| Lipase | Cortex Biochem | CR8018M1 |
| NFκB | Cell Signaling | cs-8242 |
| p-NFκB | Cell Signaling | cs-3033 |
| PERK | Cell Signaling | cs-3192 |
| pPERK | Santa Cruz | sc-32577 |
| Smooth muscle actin | Sigma-Aldrich | A7607 |
| XBP1 | Santa Cruz | sc-7160 |

#### qPCR primers

Quantitative PCR primers were designed using NCBI Primer Blast unless otherwise indicated (Table 2).

**Table 2.**
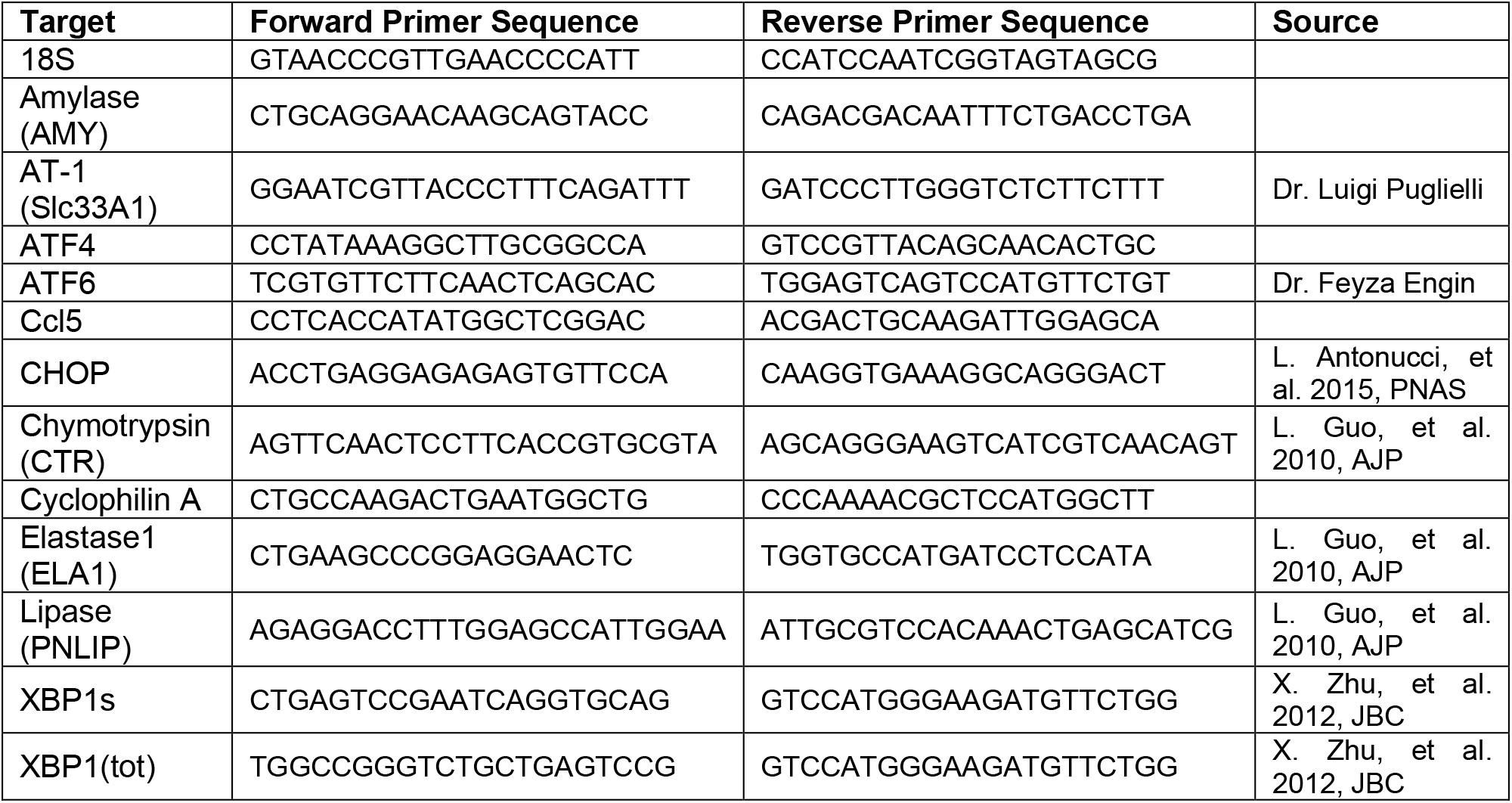
Primers for qPCR.

### Mouse Studies

#### Production of tamoxifen-inducible, acinar-specific AT-1 KO mice

Slc33a1^tm1a(KOMP)Wtsi^ heterozygous mice (C57Bl/6NTac background; Design ID: 44962; Project ID: CSD28391) were purchased from the University of California-Davis Knockout Mouse Project Repository and bred with transgenic mice expressing Flpo recombinase (CAG-Flpo1Afst/Mmucd heterozygous) to remove the targeting cassette to produce AT-1^fl/+^ mice. These mice were then crossed together and offspring without Flpo expression were used to propagate the colony. Subsequent AT-1^fl/+^ or AT-1^fl/fl^ mice were then crossed with transgenic mice expressing a tamoxifen-inducible Cre recombinase driven by the acinar-specific elastase promoter (Tg(Cela1-cre/ERT)1Lgdn/J, gifted by Drs. Craig Logsdon and Baoan Ji) (25). About 50% Cre activity in the absence of tamoxifen has been demonstrated in this elastase Cre model after 2 months; significantly, however, administration of tamoxifen induces Cre activity with 100% efficiency which remains stable for 2 years (25). The number of animals used in each experiment is shown in the figure legends. Experimental numbers may vary as not all mice were analyzed for all parameters. Only male mice were studied as we have identified inflammatory persistence in female mice with tamoxifen treatment.

Ela-Cre AT-1^fl/+^ and Ela-Cre AT-1^fl/fl^ mice at 2-3 months of age were treated with tamoxifen (MP Biomedicals, 3mg/40g BW dissolved in 98% corn oil [Sigma] and 2% ethanol [Sigma]) by oral gavage once daily for three days to induce Cre recombinase activity. AT-1^fl/+^ and AT-1^fl/fl^ mice were used as controls. Tissues were harvested at the indicated time points and analyzed as described in the sections below.

#### In vivo pancreatitis

To induce acute pancreatitis, mice were given seven intraperitoneal injections of caerulein (MP Biomedicals, 50μg/kg BW), one injection each hour for seven hours, for two consecutive days. To assess pancreatitis recovery, mice were harvested 7 days after pancreatitis induction.

### Acinar Cell Isolation and Amylase Secretion Assay

Pancreatic acini were isolated from mouse pancreatic tissue by collagenase digestion as previously described (42). Acini were incubated in salt-balanced HEPES buffer with or without CCK-8 (Research Plus) as indicated at 37°C for 30 min. Amylase released into the medium was measured using Phadebas Amylase Assay tablets (Magle Life Sciences) as previously described (43). DNA concentration was measured using Hoechst reagent (End Millipore), and secretion was expressed as amylase absorbance units normalized to total cellular DNA as done previously (44).

### Trypsin Activity Assay

Trypsin activity was measured in homogenates of pancreatic acini by a fluorogenic assay as previously reported (45). Briefly, the tissue was homogenized in a buffer containing 5 mmol/L MES, 1 mmol/L MgSO4 and 250 mmoles/L sucrose (pH 6.5). An aliquot of the homogenate (100-200 µg of protein) was incubated at 37°C for 300 sec in assay buffer containing 50 mmol/L Tris (pH 8.0), 150 mmol/L NaCl, 1 mmol/L CaCl2 and 0.1 mg/ml BSA and Boc-Gln-Ala-Arg-AMC as a specific substrate for trypsin. Cleavage of this substrate by trypsin releases 7-amino-4-methylcoumarin (AMC), which emits fluorescence at 440 nm (λem) with excitation at 380 nm (λex). Trypsin activity in each sample was determined using a standard curve for purified trypsin (Sigma Chemical).

### Plasma Amylase Assay

Plasma was prepared by centrifuging mixed arteriovenous blood from each mouse at 3,000x*g* for 15 min at 4°C. Plasma amylase activity was then determined using the Phadebas Amylase Assay tablets (Magle Life Sciences).

### Immunoblotting

Pancreatic tissue harvested from mice were homogenized in a Tris-Base buffer containing Triton-X detergent (0.2%, Sigma) and supplemented with benzamidine (1 mM, Sigma), soybean trypsin inhibitor (0.1 mg/mL, Gibco), phenylmethanesulfonylfluoride fluoride (1 mM, Sigma), and protease inhibitor cocktail (1%, Calbiochem). Tissue was homogenized using a Tissue Tearor (Dremel) followed by centrifugation at 1,000xg at 4°C for 10 min to clear cellular debris. Protein concentration of the supernatant was determined using BioRad Protein Assay Dye Reagent Concentrate. SDS-PAGE and immunoblot analyses were done as previously described(46). Membranes were stained with Ponceau S solution (Sigma) to assess relative total protein load per lane as loading controls. Antibody information is provided in Table 1.

### qPCR

Harvested pancreatic tissue was stored in RNA*later* RNA Stabilization Reagent (Qiagen) for later RNA extraction. RNA was extracted from pancreatic tissue using Qiagen RNeasy Plus Mini Kit, starting with approximately 15 mg of tissue that was homogenized in RLT buffer using a Tissue Tearor (Dremel). The concentration and purity of the RNA was assessed using Nanodrop and agarose gel analyses. RNA was then used for cDNA synthesis using the iScript cDNA synthesis kit (BioRad). cDNA templates were used for real-time quantitative PCR with KAPA SYBR Fast qPCR kit (KAPA Biosystems) and analyzed with the Roche LightCycler 480. Relative transcript levels were calculated using the 2^-∆∆Ct^ method using CycA and 18S as internal controls. Primer pairs are provided in Table 2.

### Hematoxylin and Eosin Staining

Pancreatic lobules were fixed in 4% paraformaldehyde (Ted Pella) overnight, then washed with PBS and placed in 70% ethanol (Sigma). Samples were embedded in paraffin and processed for H&E at the TRIP Laboratory facility at the University of Wisconsin.

### Picro Sirius Red and Fast Green Staining

Standard deparaffination of tissue is followed by 2 min in 0.2% phosphomolybdic acid (Fisher), brief rinsing in distilled water and then 15 min in 0.1% Fast Green (Fisher) dissolved in saturated picric acid (Fisher). Slides are then rinsed in distilled water and placed in 0.1% Fast Green dissolved in Picro Sirius Red F3BA solution (VWR) for 1 h, then rinsed in 0.1 N HCl for 2 min, followed by brief rinses in 70%, 95%, 100% ethanol and finally 5 min in xylene (Fisher) (in that order). Slides are then mounted with coverslips using solvent based mounting media.

### High Pressure Freezing Electron Microscopy

*In vivo* animal fixation was used according to approved animal protocol. Animals were anesthetized; a mixed solution of 2% formaldehyde (Ted Pella) and 2% glutaraldehyde (Electron Microscopy Sciences) in 1X Sorensen’s PB (Electron Microscopy Sciences) was allowed to perfuse briefly into the animals’ circulatory system via the heart. Next the pancreas was removed, trimmed and allowed to fix overnight in 2% formaldehyde/2% glutaraldehyde. Tissue was then dissected into 1mm X 1mm pieces, rinsed well with 0.1 M PB, dipped in 20% BSA in filtered water, placed on planchets coated with hexadecane (Ted Pella) and introduced into the HPF machine. Samples next underwent freeze substitution and dehydration over 2 days in 2% osmium tetroxide (Electron Microscopy Sciences) and acetone (Fisher) with liquid nitrogen. Samples were then embedded in Epon, sectioned and processed as previously described for imaging (47).

### Immunofluorescence Microscopy

Immunofluorescence microscopy was conducted on cryostat sections of 4% paraformaldehyde fixed pancreatic lobules as previously described (43). Blocking and incubations were done in PBS supplemented with 3% bovine serum albumin (Calbiochem), 2% goat serum (Sigma), 0.7% cold-water fish skin gelatin (Sigma), and 0.2% Triton X-100 (Sigma). Cryostat sections were rinsed with PBS then treated with Image-iT FX (Invitrogen) according to manufacturer instructions. Primary antibodies were added simultaneously for 2 hrs at RT. Sections were washed with PBS then incubated with secondary antibodies for 1 hr at RT. If indicated, tissue was incubated with Alexa-conjugated phalloidin (Invitrogen) to label actin according to manufacturer instructions. Sections were washed again with PBS and mounted with coverslips and ProLong Gold antifade reagent with DAPI (Invitrogen) to label nuclei.

### Statistics

All data are expressed as mean ± SEM. Calculations were performed using GraphPad Prism 8.0. Two-tailed unpaired Student’s *t* test was used for comparison between 2 groups; F-tests were performed prior to *t* test analysis to determine equality of variance. One-way or two-way ANOVA was used for multiple group comparisons. Statistical significance levels are indicated as *p<0.05, **p<0.01, ***p<0.001, ****p<0.0001.

### Study Approval

The University of Wisconsin Committee on the Use and Care of Animals approved all studies involving animals.

## Author Contributions

GEG and LP conceived and directed the study. GEG, MMC, AL, SJP, and LP designed and interpreted the experiments. Experiments were performed and data analyzed by MMC, DDHT, AL, KD, and YP. Figures were prepared by MMC, DDHT, and KD. GEG and MMC wrote the manuscript in consultation with all authors.

## Acknowledgements

This work was supported by the following grants: NIH RO1DK07088 (GEG), NIH PO1DK098108 (GEG, AL, and SJP), NIH RO1NS094154 (LP), NIH RO1AG053937 (LP), NIH RO1AG057408 (LP), NIH T32DK007665 (MMC), and USDA/HATCH WIS01583 (GEG).

---

**Supplementary Figure 1.**
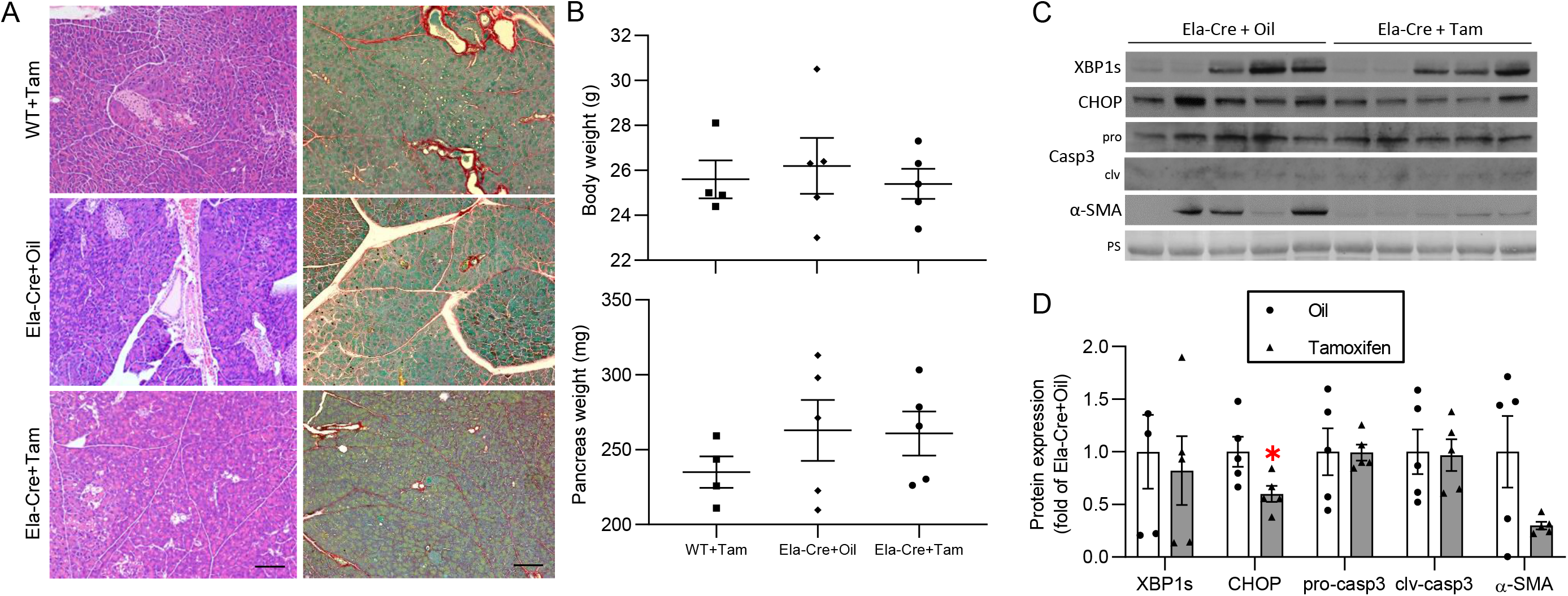
**A.** Representative H&E and Sirius red staining of pancreas from WT or Ela-Cre mice given oil or tamoxifen (Tam), as indicated. Scale bar: 50μm. **B.** Body weight and pancreatic weight. **C.** Immunoblot of XBP1s, CHOP, caspase3, and α-SMA, quantified in **D.** Ponceau S (PS) is loading control. Data are mean +/− SE, n=5, *p<0.05. Statistical comparison of means was done by unpaired two-tailed Student’s *t*-test.

